# CRISPR/Cas “non-target” sites inhibit on-target cutting rates

**DOI:** 10.1101/2020.06.12.147827

**Authors:** Eirik A. Moreb, Mitchell Hutmacher, Michael D. Lynch

**Affiliations:** Department of Biomedical Engineering, Duke University, Durham, NC

**Keywords:** Cas9, cutting rates, specificity, non-target pool

## Abstract

CRISPR/Cas systems have become ubiquitous for genome editing in eukaryotic as well as bacterial systems. Cas9 associated with a guide RNA (gRNA) searches DNA for a matching sequence (target site) next to a protospacer adjacent motif (PAM) and once found, cuts the DNA. The number of PAM sites in the genome are effectively a non-target pool of inhibitory substrates, competing with the target site for the Cas9/gRNA complex. We demonstrate that increasing the number of non-target sites for a given gRNA reduces on-target activity in a dose dependent manner. Furthermore, we show that the use of Cas9 mutants with increased PAM specificity towards a smaller subset of PAMs (or smaller pool of competitive substrates) improves cutting rates. Decreasing the non-target pool by increasing PAM specificity provides a path towards improving on-target activity for slower high fidelity Cas9 variants. These results demonstrate the importance of competitive non-target sites on Cas9 activity and, in part, may help to explain sequence and context dependent activities of gRNAs. Engineering improved PAM specificity to reduce the competitive non-target pool offers an alternative strategy to engineer Cas9 variants with increased specificity and maintained on-target activity.

**Highlights:**

- The pool of non-target PAM sites inhibit Cas9/gRNA on-target activity
- non-target PAM inhibition is dose dependent
- non-target PAM inhibition is a function of gRNA sequence
- non-target PAM inhibition is a function of Cas9 levels

## Introduction

CRISPR/Cas systems were first harnessed in 2012 for DNA editing and have provided numerous exciting advances and future opportunities in biotechnology.^1^ These systems enable RNA targeted cutting and/or binding to DNA in a sequence specific manner, wherein a Cas enzyme binds with a targeting RNA to form a Cas/gRNA complex, which can specifically recognize a DNA target.^2^ Not only have these systems been rapidly deployed as research tools in studying all domains of life from bacteria to humans, but they have offered new routes to advanced therapies for many diseases, from engineered cell lines such as CAR-T cells to editing humans genomes, offering the potential of cures for numerous genetic diseases.^3–6^ CRISPR/Cas systems and *S. pyogenes* (Cas9) in particular (the most well studied) work remarkably well across a diversity of species and applications and are easier to use than the tools previously available.^7^ However, challenges remain in the use of CRISPR/Cas systems, even with *S. pyogenes* Cas9. These include the risk of unwanted off-target DNA activity and gRNA- and context-dependent (host organism, cell type, etc..) differences in cutting rates. These remaining challenges highlight basic gaps in our understanding of factors affecting CRISPR/Cas activity.

This basic challenge underscores gaps in our understanding of factors affecting CRISPR/Cas activity. While simple over-expression of Cas9 can be used to improve on-target activity, it is also linked with increased off-target effects.^8^ These challenges are amplified in certain circumstances, including delivery of Cas9 as mRNA or ribonucleoprotein (RNP) where low Cas9 expression and short Cas9/gRNA half-life limit on-target activity,^9^ or using high fidelity versions of Cas9 that have been shown to sacrifice on-target activity for specificity.^9–15^ Broadly speaking, many of the strategies to reduce off-target activity result in lower on-target activity, either through reduced Cas9 expression or by sacrificing on-target activity for higher fidelity. Additionally, numerous studies have demonstrated the Cas9 activity is not only dependent on a specific gRNA sequence but also on the genomic context (host organism, cell type, etc.).^16,17^ The same gRNAs can have highly variable activity between animal models, cell lines, and even strains of *E. coli*. ^16–18^ In this study, we confirm that “non-target” sites, representing a significant level of context dependent variability, competitively bind to Cas9/gRNA complexes, and can significantly impact on-target activity of Cas9/gRNA complexes *in vivo*.

While DNA cleavage occurs at “on-target” and “off-target” sites (with sequences similar to a given “on-target” site), “non-target” sites make up a diverse group of PAM containing loci where Cas9/gRNA complexes can bind, but where DNA cutting does not occur. Previous studies have demonstrated that the time Cas9/gRNA complexes spend interrogating any “non-target” site is dependent on the sequence similarity between this site and a given gRNA and can range from 0.1 to over 100 minutes per site.^19^ In this sense, the collection of non-target sites in a genome represents a large pool of potential competitive inhibitory substrates that is dependent both on Cas9 expression level and gRNA sequence. As illustrated in Figure 1, the pool of non-target PAM sites make up a vast potential sequence space, through which the Cas9/gRNA complex must search for its target. Cas9 interrogates each PAM in this non-target pool or “search space”, in a sequential, zipper-like fashion where the PAM-proximal sequence is checked for a match first (Figure 1a). The search space is dependent on the genomic context and is a variable which is orthogonal to the enzymatic activity of Cas9 or the target dependent activity of any given gRNA (Figure 1e-g).

**Figure 1:**
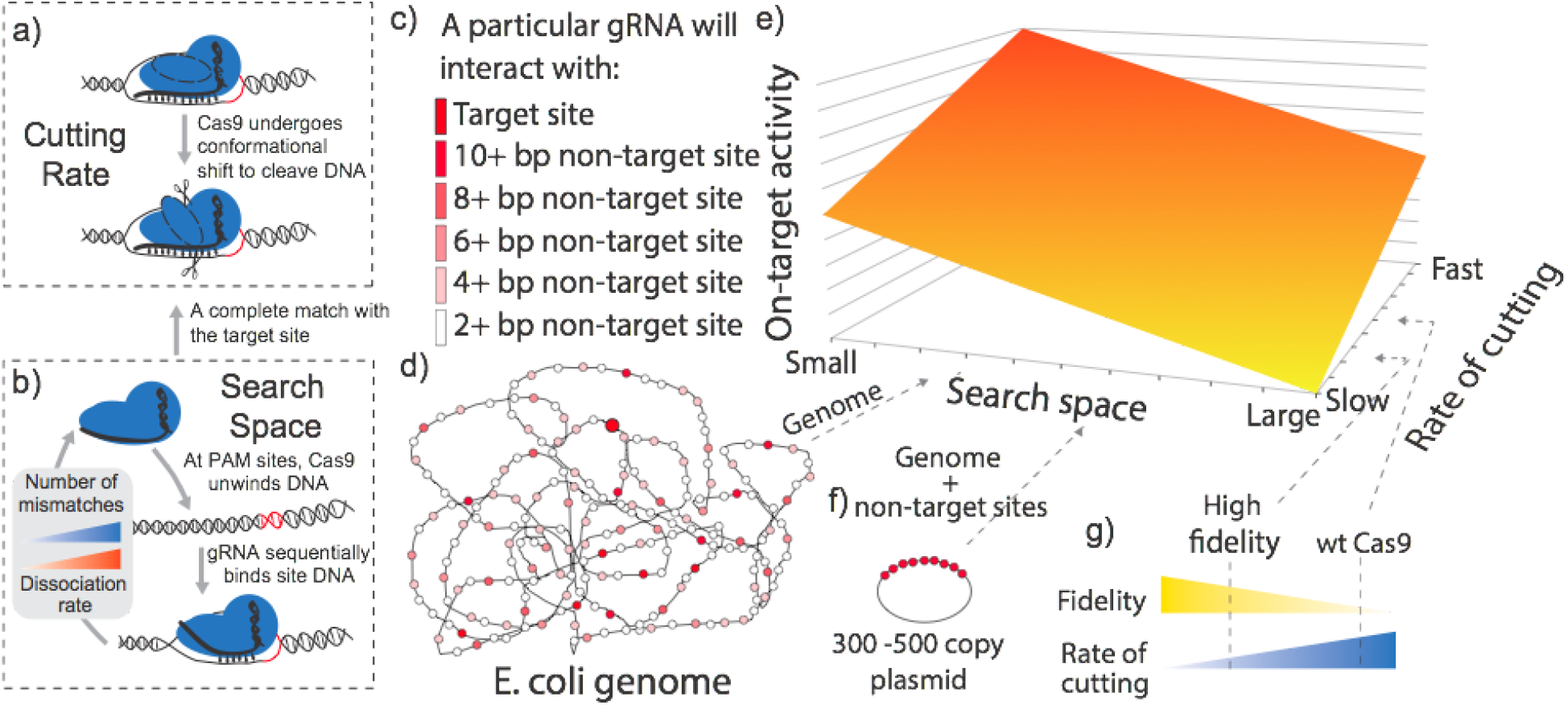
Cas9 activity is dictated by its ability to a) cut DNA but also how it b) finds the target. c-d) A Cas9/gRNA complex will interact with numerous PAM sites around the chromosome with different degrees of sequence identity compared to its target. e) We define the non-target pool or search space as an orthogonal axis to Cas9 cutting. f) the non-target pool can be modified by the inclusion of extrachromosomal DNA. g) Cas9 mutations are known to affect cutting rate. For example higher fidelity Cas9 mutants have decreased cutting rates when compared to wild type Cas9.

## Methods

### Media & Reagents

Low salt Luria broth (LB Lennox, VWR #90003-114) was used for cloning, strain propagations, and experiments in both liquid cultures and agar plates. Agar A was used for agar plates (BioBasic #9002-18-0). Antibiotics were obtained from Sigma-Aldrich (St. Louis, MO). Antibiotic concentrations, unless otherwise stated, were as follows: chloramphenicol (35 μg/mL), spectinomycin (30 μg/mL), kanamycin (35 μg/mL). Anhydrotetracycline (aTc; used to induce Cas9 and gRNA expression) was prepared in filtered 70% ethanol at a 1000× stock and used at a final working concentration of 100 ng/L.

### Strains & Plasmids

Strain E. cloni (Lucigen, Middleton WI) was used for cloning and plasmid propagation. Strain BW25113, used for all killing assays in Figures 1–5, was obtained from the Yale E. Coli Genetic Stock Center (http://cgsc.biology.yale.edu). Strain W was used for the library assay in Figure 6 and obtained from ATCC (ATCC #9637). Guide RNA plasmids were constructed using “around the world” (ATW) PCR followed by kinase, ligase and DpnI treatment (KLD reactions), using the pKDsgRNA-non-targeting plasmid as a template. Cas9 variant plasmids were constructed via ATW PCR and a KLD reaction and/or via DNA assembly using New England Biolabs (NEB) HiFi DNA Assembly (NEB E5520S). Refer to Table S2 for more details on cloning methodologies for each plasmid. pNT-15 was constructed using NEB HiFi DNA Assembly with a gBlock® from IDT (Coralville, IA) and cloning vector pSMART-HCKan (Lucigen, #40704-2). All PCR reactions were performed with Q5® Hot-Start High Fidelity Master Mix (NEB, M0493S). Reagents for the KLD reactions came from NEB: T4 PNK (M0236S), T4 DNA ligase (M0202S), and DpnI (R0176S). All primers and sequences used in construction of these plasmids are in Supplemental Table S2 along with descriptions. All plasmid sequences were confirmed via Sanger Sequencing at Genewiz (Morrisville, NC).

### Survival Assays

Survival assays were performed as previously described.^16^ Cells were first transformed with the gRNA plasmid (pKDsgRNA-xx, where xx indicates the gene it targets) and recovered at 30°C. The origin of replication is temperature sensitive so all subsequent experiments were performed at 30°C. To test individual gRNA activity, cells were made electrocompetent again and transformed with one of the Cas9 plasmids. Cells were then recovered for 2 hours in 300uL of LB and plated in serial dilutions on plates with and without aTc and both containing spectinomycin and chloramphenicol. Colony forming units (CFUs) were counted on each plate and adjusted for dilution. The survival frequency was then calculated as the number of colonies on the aTc plate (Cas9 induced) divided by the number of colonies on the plate without aTc (no Cas9 expression). Survival frequencies were then averaged over three transformations. For assays with the non-target plasmids (pNT-x), the assay was performed similarly except that the cells contained the gRNA plasmid and non-target plasmid prior to electroporation with Cas9 plasmids. In addition, kanamycin was added to the plates to select for the non-target plasmids.

### gRNA Library Design

The gRNA library was designed as a subset of the library previously reported by Moreb et al. and constructed using the same methodology.^16^ The original library was designed to have a unique 12 bp seed region, be a minimum of 320bp from other gRNA targets selected and be present in both BW25113 and MG1655. In this library, we selected 8,200 gRNA from the previous library that also targeted strain W. A BsmBI site, overhang (underlined), was added onto the ends of each spacer sequence, and primer binding sites were added as follows:
5′-AGGCACTTGCTCGTACGACGCGTCTCA<u>GCAC</u>-[20 bp spacer sequence]-<u>GTTT</u>CGAG-ACGATGTGGGCCCGGCACCTTAA-3′.

These guide sequences were ordered as a single library of 8,200 oligos from Twist Biosciences (San Francisco, CA). The pooled library was PCR amplified using EAM026 and EAM027. pKDsgRNA-non-targeting was designed as both a nontargeting control and a cloning vector containing BsmBI sites to insert the gRNA library. Golden Gate Assembly was performed to ligate the amplified gRNA pool into the pKDsgRNA-non-targeting backbone. Briefly, Esp3I (Thermo Fisher Scientific Cat# ER0451) was used in the cleavage step while T4 Ligase (NEB M0202S) was used for ligation. The library was then transformed into Lucigen’s 10G Elite electrocompetent *E. coli.* The Carbon-Clarke equation was used to calculate how many colonies were needed to ensure a 99% confidence that all guides in the library was represented after each transformation (∼37,000 colonies).^20^ Colonies were scraped and midi-prepped (Zyppy Plasmid Midiprep Kit, Zymo Research, Irvine, CA) in preparation to be transformed into strain W.

### gRNA Library Selection Assay

Strain W was transformed with the gRNA library, again ensuring enough colonies were collected to ensure the whole library was present (>37,000 CFUs). Colonies were scraped from plates and inoculated into LB to make electrocompetent cells, which were then transformed with either pEM-Cas9-HF1 or pEM-Cas9-HF1_D1135E_. Cells were recovered for two hours in LB without antibiotics and then spun down and resuspended at 1 OD in two cultures of 800 uL LB. Both cultures contained chloramphenicol and spectinomycin to maintain selection. The cells were then cultured in deep 96-well plates for 8 hours at 30°C, 300rpm. Plasmids were then purified (Zyppy Plasmid Miniprep Kit, Zymo Research, Irvine, CA) and PCR of the gRNA sequence was performed using custom barcoded PCR primers (Supplemental Table S2, primers EAM032-EAM047). Sequencing was performed on an Illumina HiSeq 4000 at Duke Sequencing and Genomic Technologies Shared Resource using 150bp paired end reads. Read and index primers are EAM028-031 in Supplemental Table S2. Reads were de-multiplexed using deML and counts tables were generated using bowtie2 and samtools, as described previously.^16,21–23^ For each barcoded sample, read counts were converted to frequencies by dividing each read by the total number of reads. The log2 of the fold change between samples with aTc and without aTc was then calculated. gRNA with 0 reads in either the aTc or no aTc sample were thrown out and the remaining samples were averaged across four replicates (n=2, 3, or 4, depending on the number of samples with enough reads for an individual gRNA). In order to compare Cas9-HF1 and Cas9-HF1_D1135E_, we selected the 6,044 gRNA with averaged reads for both Cas9 variants.

## Results

### Measuring “on target” cutting rates via an E. coli survival assay

To test the impact of the non-target pool on Cas9 activity, we first needed an assay to rapidly measure Cas9/gRNA complex cutting rates and turned to *E. coli* as a model system. As an easy-to-handle model organism, *E. coli* has been used for the development of high-fidelity Cas9s, ^12^ to successfully compare on-target activity of wild type Cas9 with eCas9 ^24^ and to engineer new PAM specificities. ^25^ In *E. coli*, double stranded DNA (dsDNA) breaks can lead to death and CRISPR/Cas systems have been used for the selection of genetically modified strains.^16^ dsDNA breaks lead to induction of the SOS response, which results in increased RecA (a rad51 homologue) based homologous recombination. ^26^ As a typical *E. coli* cell contains multiple chromosomes (from 1 to 8, depending on growth rate), ^27^ sister chromatids offer templates for recombination based repair of Cas9 induced cuts. Thus, as illustrated in Figure 2a, Cas9-mediated cell death takes place only if all copies of the chromosome are cleaved faster than they are repaired. ^16^ A benefit of this experimental system, over many eukaryotic systems, is that for efficient killing to occur the rate of CRISPR/Cas cutting has to exceed the rate of repair. As a result, cell death due to DNA cleavage is limited *in vivo* by DNA cutting rates and killing efficiency can therefore be correlated to CRISPR/Cas activity.^16^ This experimental system enables us to measure survival frequency or killing efficiency (Figure 2b) by simply counting colonies.

**Figure 2:**
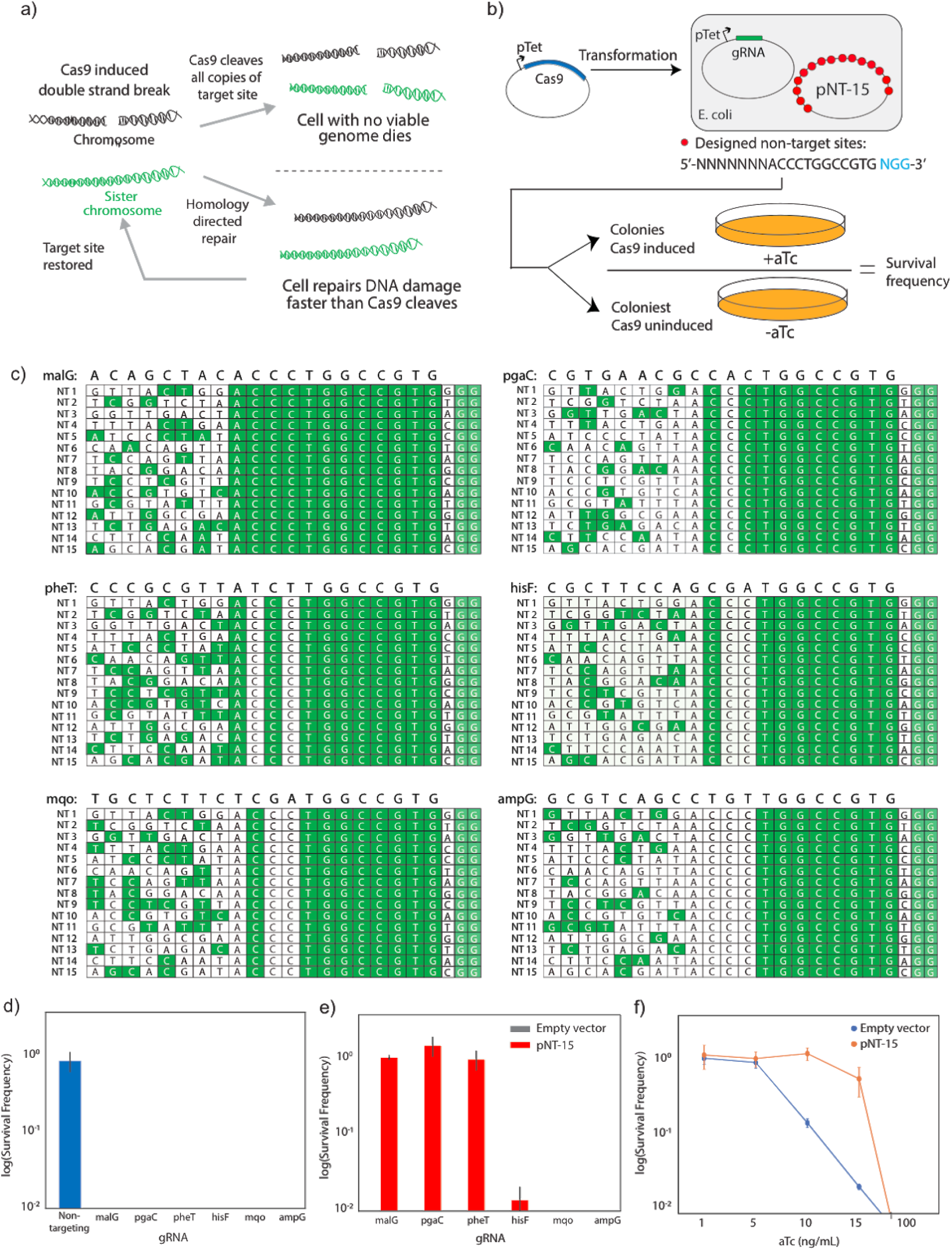
Cas9 on-target activity can be inhibited by the addition of plasmid based partially matching non-target sequences. a) In *E. coli*, Cas9 activity is a competition between Cas9 mediated double strand breaks (DSB) and repair of the cleaved sites, where efficient Cas9 cleavage leads to cell death but inefficient cleavage allows the cell to repair its genome and survive.. b) Using an assay in which we artificially increase the number of non-targets (pNT-15), we evaluate Cas9 activity by calculating counting colonies on agar plates with and without aTc, which induces Cas9 expression. The number of colonies that survive Cas9/gRNA expression over the number of colonies without Cas9 expressed gives the survival frequency. c) The pNT-15 plasmid contains 15 non-target sites with the same PAM proximal 12 bp. Shown are alignments with six gRNA to each of the non-target sequences. Longer sequence matches would be expected to be stronger competitive inhibitors. d) Each of the six gRNA evaluated efficiently kills *E. coli* e) However, with the addition of pNT-15, malG, pgaC, and pheT are completely inhibited. HisF shows incomplete inhibition, while mqo and ampG don’t appear inhibited. f) Evaluating mqo activity with decreasing Cas9 expression (induction by aTc).

Using this assay, we measured baseline killing efficiencies of 6 gRNAs, targeting loci in 6 different ORFs including: *ampG*, *hisF*, *malG*, *mqo*, *pgaC*, and *pheT* (hereafter referred to as such) where Cas9 is expressed under the control of the inducer anhydrotetracycline (aTc). Results are given in Figure 2d. All 6 gRNA led to complete killing or 0% survival at least within resolution of the assay. Refer to Supplemental Tables for a list of strains, plasmids and DNA used in this study, and the Methods Section for experimental details.

### Increasing the non-target pool decreases on-target activity

Having confirmed the baseline killing activity for a set of gRNAs, we next set out to show that expanding the pool of non-target sites could reduce on-target activity in *E. coli*. To increase the non-target pool, we designed 15 non-target sites with PAM-proximal sequences partially matching the 6 gRNAs (Figure 2c). While all non-target PAM sites can be thought of as competitive inhibitors, as discussed above, those with more sequence similarity (Figure 2c) would be expected to be stronger inhibitors. These non-target sites were cloned into a plasmid (pNT-15, Fig 2b) with an estimated copy number of 400/cell effectively creating strains with an altered competitive PAM pool for each gRNA. We then measured the killing efficiency of each of these 6 gRNA in cells bearing plasmid pNT-15. Results are given in Figure 2e. These non-target DNA sequences reduced the killing efficiency of 4 of the 6 gRNAs to varying degrees. The reduction in killing was qualitatively a function of the sequence similarity of non targets to a given gRNA (Figure 2c). For the mqo targeting gRNA, the increase in the non-target pool provided by pNT-15 was insufficient to inhibit killing. The ability of Cas9/gRNA complexes to find and cleave a particular site in a manner that leads to cell death is in part based on availability of sufficient Cas9/gRNA complexes. We expected that as the Cas9/gRNA concentration decreases, we may see greater inhibition. Indeed, as we decreased Cas9 expression by lower aTc (inducer) levels, we show that killing by mqo gRNA is inhibited (Figure 2f). We next sought to assess whether the observed level of inhibition was dose dependent. We constructed derivatives of plasmid pNT-15 with fewer non-target sites and saw a dose dependent effect, as illustrated in Figure 3. These results confirm that increasing the non-target pool decreases on-target activity and that competitive binding to non-targets sites can impact CRISPR/Cas on-target activity. Additionally, we have a simple and titratable assay to measure the impact of non-target inhibitors.

**Figure 3:**
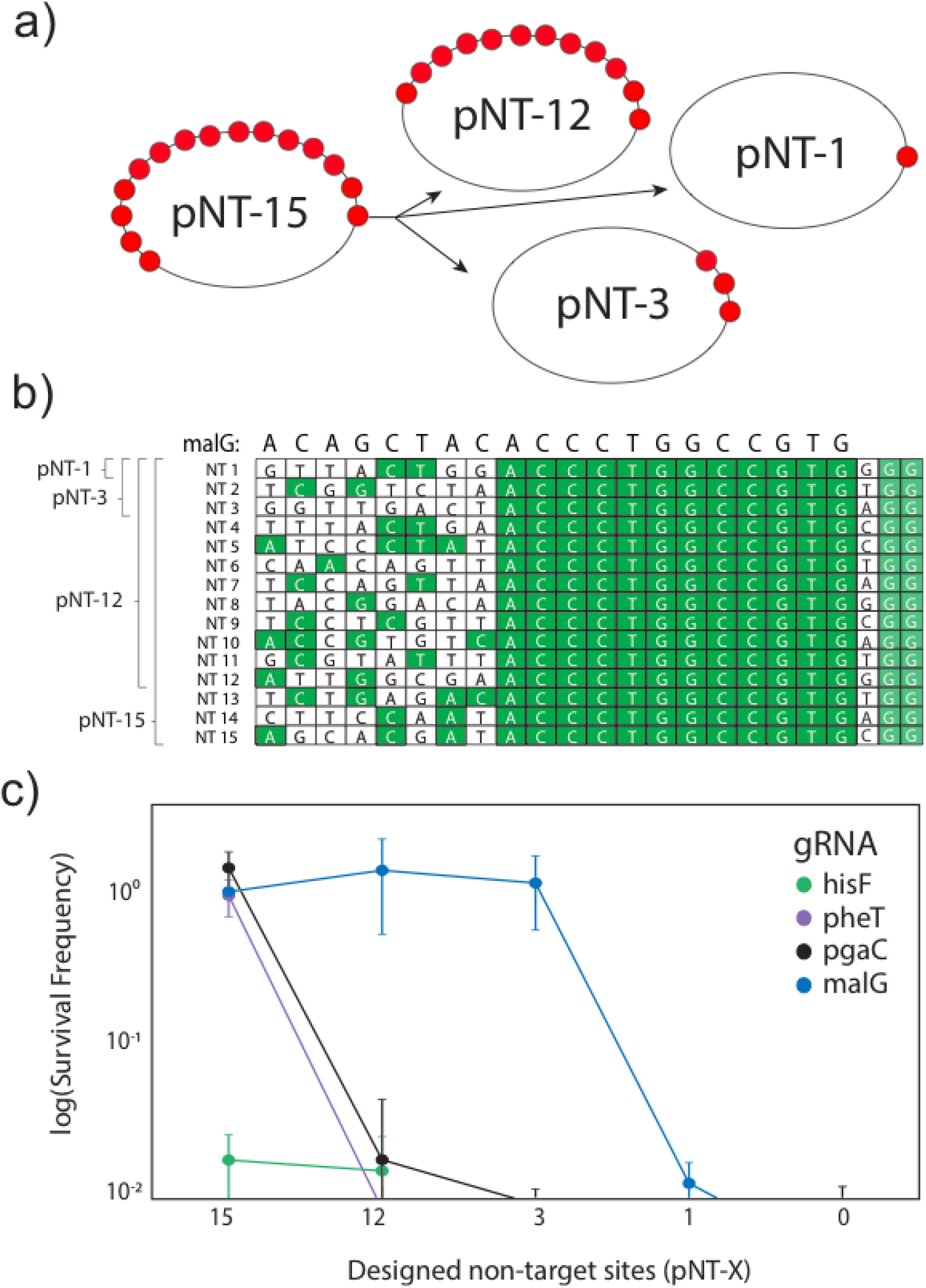
Dose dependence of non-target inhibitory PAMs. a) Construction of plasmids containing different numbers of non-target sequences where the exact sequence of the non-target sites for each plasmid is indicated in b) (showing the alignment with malG gRNA). c) Decreasing the number of non-target sites shows a dose-dependent response of inhibitory non-targets on the survival frequency of cells expressing various gRNAs (the X in pNT-X refers to either 15, 12, 3 or 1 along the X axis).

### Decreasing non-target pool inhibition increases on-target activity

In order to better understand the impact of the larger non-target pool on Cas9 activity we took a complementary approach to the one described above and focused on decreasing the non-target pool, which would be expected to increase on-target activity. To accomplish this, we chose to use a Cas9 variant with increased PAM specificity. Reducing the number of PAM sites recognized by Cas9 effectively reduces the size of the inhibitory non-target pool. Or more accurately reducing the affinity for additional PAMs will reduce the level of competitive inhibition of these non-targets. Wild type Cas9 recognizes the canonical PAM site “NGG” but has also been shown to bind to non-canonical PAM sites, most prominent of which are “NAG” and “NGA” but with reports of activity at several others as well.^25^ In *E. coli,* there are ∼540,000 “NGG” sites and additional ∼470,000 “NAG” and ∼530,000 “NGA” non-canonical PAMs, representing a total non-target pool of over 1.5 million PAM sites as shown in Figure 4. A mutant, Cas9_D1135E_, has previously been reported to have much higher specificity for the canonical “NGG” PAM site, potentially reducing the non-target pool by up to ⅔ (∼ 1 million sites).^25^ Interestingly, this mutant has also been shown to reduce off-target activity.^25,28^ We hypothesized this increased specificity is due to reduced binding to all PAMs, thereby limiting the affinity for all sites without complete gRNA matches. This would effectively reduce the potency of the inhibition of the non-target pool by reducing the affinity for non-targets with NGG PAMs as well as non-targets with non-canonical PAMs. We therefore compared the killing efficiencies of Cas9 and Cas9_D1135E_ with the malG, pgaC and pheT gRNAs. As killing efficiencies of these gRNAs in wild type *E. coli* were very high (Figure 2), we would not expect to measure any improvements. As a result, we evaluated killing efficiencies with the non-target plasmid pNT-15, where baseline killing efficiencies are reduced (Figure 2) and potential improvements could be measured. While in the presence of pNT-15, the pheT and pgaC gRNA did not lead to cell death with wild type Cas9 but did efficiently kill cells when used with Cas9_D1135E_, indicating higher on-target activity. In addition, Cas9_D1135E_ improved killing with the hisF gRNA. However, Cas9_D1135E_ did not lead to any improvement in killing with the malG gRNA. In the case of malG, the plasmid based non-target sites, with higher sequence similarity leading to more stable Cas9 binding, are very strong competitive inhibitors.

**Figure 4:**
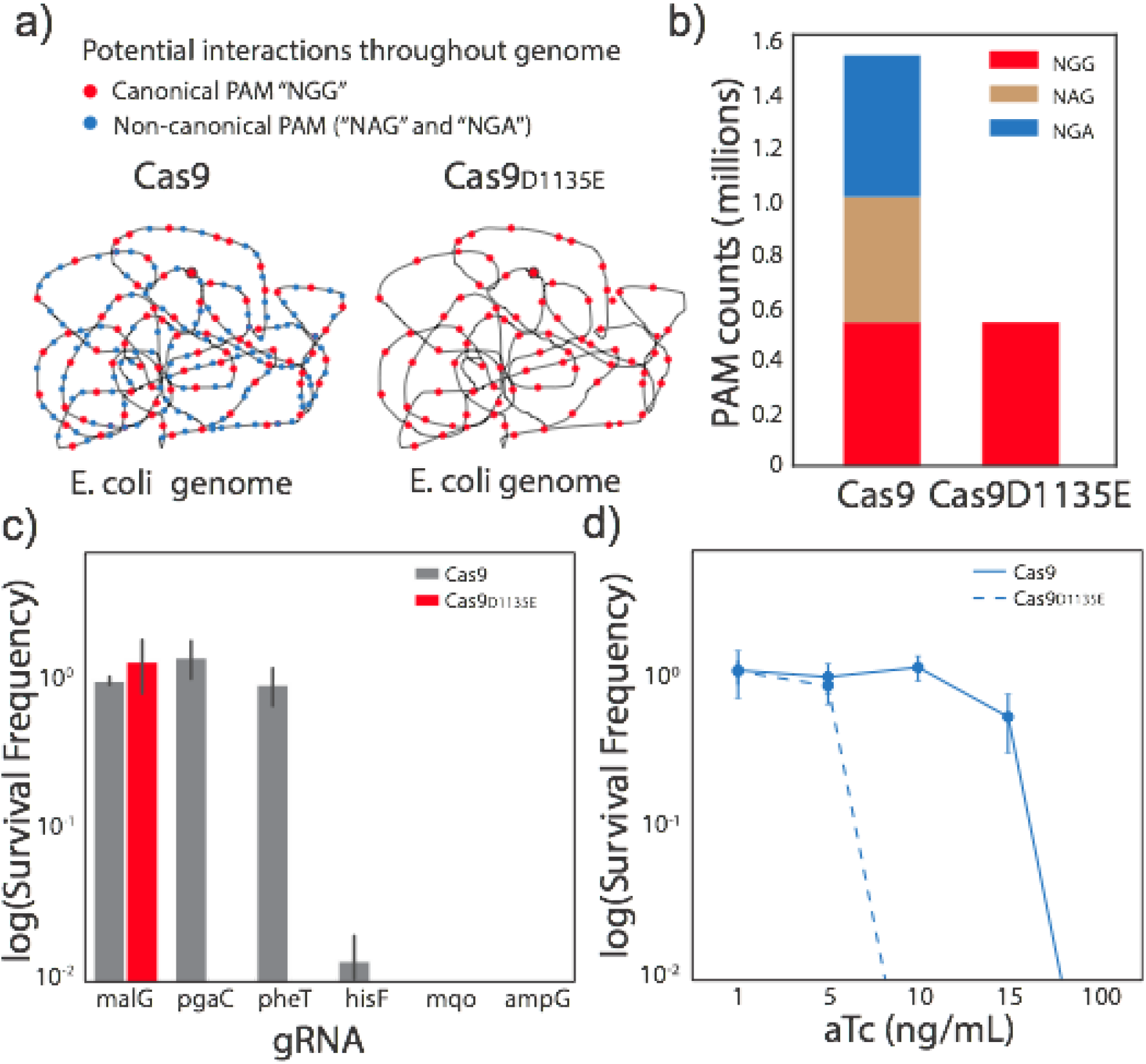
Cas9_D1135E_ reduces the non-target pool and leads to improved Cas9 on-target activity in the presence of artificial non-targets. a) Cas9 is known to recognize the PAM sequence NGG but to a lesser extent also recognizes NAG and NGA. Cas9_D1135E_ is a mutant of Cas9 that is reported to be more specific to NGG PAM sites. b) Potential reduction in the non-target pool from ∼1.5 million to ∼0.5 million, with the D1135E mutant.. c) Although Cas9_D1135E_ binds the artificial non-targets on pNT-15, it has improved on-target activity in the presence of pNT-15 compared to the wild type Cas9 for pgaC, pheT and hisF. d) Cas9_D1135E_ shows higher on-target activity at lower Cas9 expression levels when targeting mqo.

Importantly, the non-target sites on plasmid pNT-15 are designed with “NGG” PAM sites and therefore are still high affinity non-target sites for Cas9_D1135E_. As a result, differences in on-target activity between Cas9 and Cas9_D1135E_, as seen in Figure 4c, can at least partially be attributed to reducing the number or strength of non-target site inhibitors throughout the genome rather than reducing the number of sites on the pNT plasmids, suggesting that Cas9_D1135E_ improves on-target activity by reducing the number or strength of competitive inhibitors throughout the genome rather than changing interactions at the artificial non-targets. Taken together, the results in Figures 2–4, support the hypothesis that the non-target pool can significantly inhibit Cas9 on-target activity, and that the inhibition is the cumulative effect of a large number of non-target sites. The inhibition is dependent on the gRNA sequence and genomic context.

### Reducing the non-target pool increases on-target activity with slower Cas9 variants

We next sought to evaluate the applicability of reducing the non-target pool to improve the on-target activity for slower Cas9 variants, including the high fidelity variants Cas9-HF1, eCas9, HypaCas9 and OptiCas9 (Figure 5).^10,11,13,14^ These higher fidelity Cas9 variants have been engineered to have reduced off-target activity (reduced cutting at sites other than the target site) and improved on-target specificity. Mutations that lead to this increased specificity (Figure 5a), result in enzymes that take longer to cut DNA (Figure 1b), leading to more time for the Cas9/gRNA complex to leave a given site if there is not a perfect sequence match with the gRNA. Although more specific, these variants suffer from reduced cutting rates when compared to wild type Cas9.^9,10,24,29^ Using the same set of gRNAs and pNT-15, we compared the killing efficiency of wild type and high fidelity Cas9 variants with and without the D1135E mutation. Results are given in Figure 5. As expected, the high fidelity variants had reduced killing efficiency or increased survival compared to wild type Cas9. This is consistent with slower on target activities reported by others.^9–15^ The addition of the D1135E mutation resulted in improved on-target killing with all gRNAs tested. Again, the killing efficiencies were dependent on Cas9 expression levels as seen in Figures 5e-h, and the D1135E mutation resulted in increased killing even at higher expression levels.

**Figure 5:**
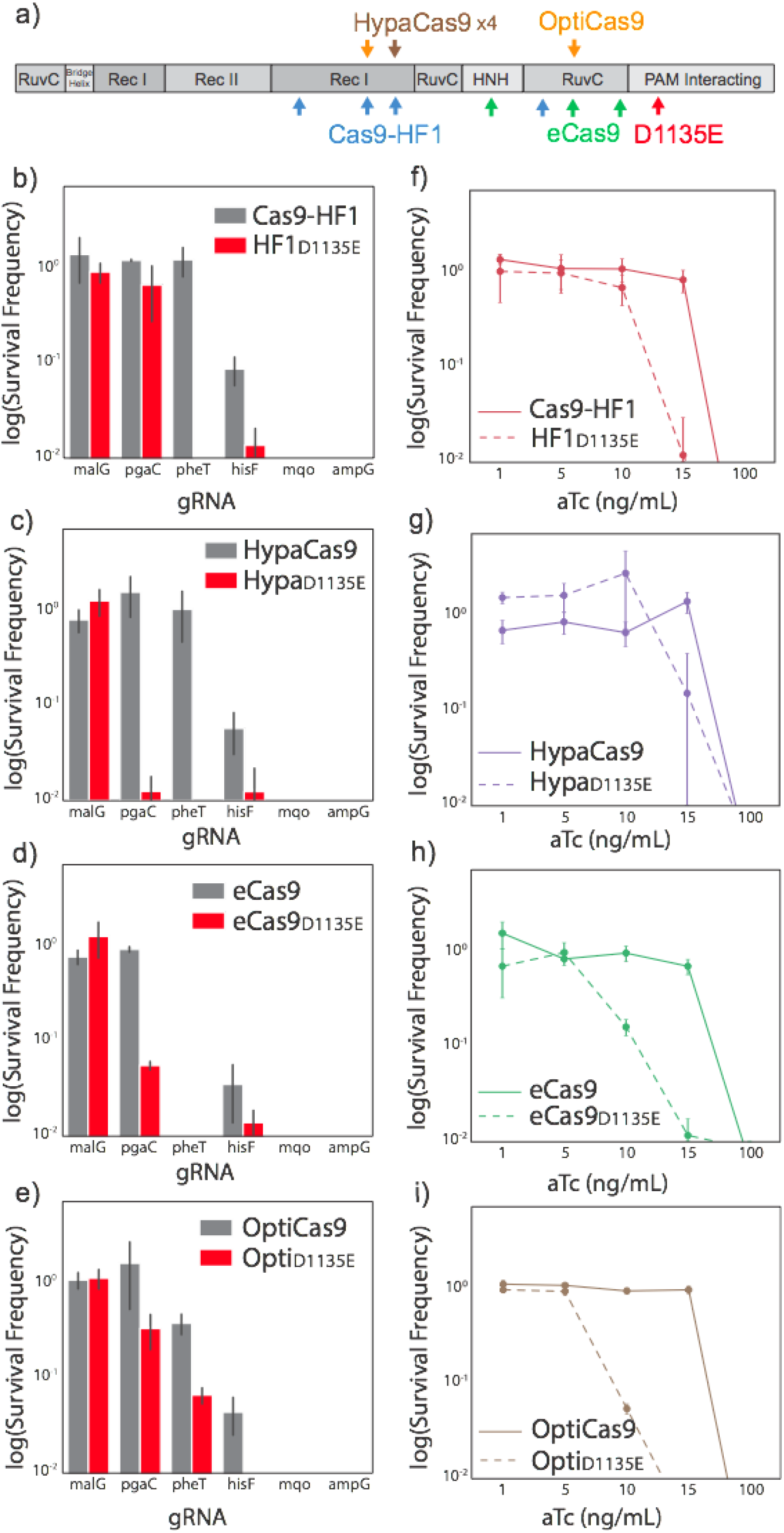
(a) A schematic of high fidelity mutations and the D1135E mutation in the primary structure of Cas9. (b-i) D1135E decreases survival (improved killing) with different gRNA (b-e) and at different Cas9 expression levels (f-i) The following high fidelity mutants were used: Cas9-HF1 (a & e), HypaCas9 (b & f), eCas9 (c & g), and OptiCas9 (d & h).

### The impact of the non-target pool is physiologically meaningful

While the results above support that non-target pool inhibition is real and measurable, they were obtained in artificially engineered systems. The question remained whether or not natural non-target pools have a physiologically meaningful impact *in vivo.* Therefore, we evaluated the activity of a genome wide library of 6,044 gRNAs in strains expressing the higher fidelity HF1 Cas9 variant with and without D1135E, the results of which are given in Figure 6. The D1135E mutation led to increased killing for a large number of gRNA sequences. Specifically, while 3,213 gRNA efficiently killed with both Cas9-HF1 and HF1-D1135E, another 1,140 gRNA only efficiently killed with HF1-D1135E compared to 94 gRNA that only worked in Cas9-HF1 (Fig. 6b). Taken together these results support that reducing the non-target pool increases on-target activity with slower Cas9 variants and that the inhibitory impact of the non-target PAM pool can be physiologically meaningful.

**Figure 6:**
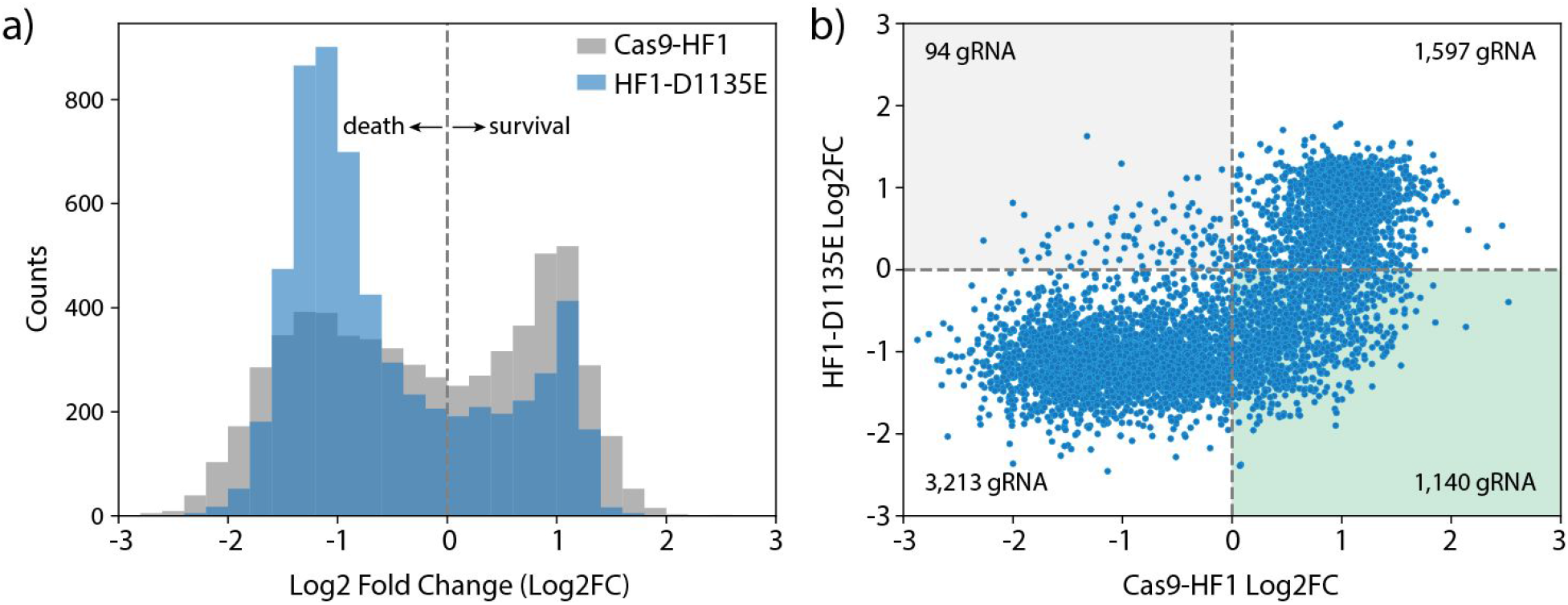
A library of 6,044 gRNA was used to evaluate (via library enrichment) Cas9-HF1 and Cas9-HF1_D1135E_ on-target activity. (a) On a population-wide level, D1135E improves on-target activity (negative Log2 fold change) for many gRNA compared to Cas9-HF1). (b) On-target activity of Cas9-HF1 against HF1-D1135E shows 1,140 gRNA with improved on-target activity using HF1-D1135E compared to Cas9-HF1 (green shaded region). Only 94 gRNA show the opposite trend (grey shaded region).

## Discussion

We have demonstrated that the non-target pool or “search space” can impact on-target Cas9/gRNA complex activity. Specifically, increasing the search space reduces on-target activity and decreasing the search space increases on-target activity. Additionally, mutations in Cas9 that reduce the search space can be used to increase on-target activity of slower high fidelity variants. As illustrated in Figure 1, the pool of non-target sites or search space represents variation orthogonal to Cas9 mutations that affect activity in a gRNA specific and context specific manner. Together these results provide a framework for both understanding and improving Cas9 on-target activity.

Numerous studies have demonstrated the Cas9 activity is not only dependent on a specific gRNA sequence but also on the genomic context. One of the largest sources of variation in these systems is the pool of non-target sites. By its very nature, the variability in Cas9 activity due to context is extremely difficult to study. To date numerous studies have in effect sampled this context by testing different species, strains, cell lines etc,. While several studies have investigated the impact of additional “off-target” in some cases non-target sites, ^30,31^ this is difficult to do in a systematic way as changes to the chromosomal context are confounded with numerous other variables in addition to the non-target pool. The importance of the “non-target” pool is difficult to assess in many experimental systems and varies depending on the gRNA utilized. Cas9 expression levels alone are a confounding variable in many studies. All of the individual guides reported in this study have 8 bp in common so share a very similar search space, yet have dramatic differences in on target activity as a function of changes in the non-target pool indicating that even current computation approaches are likely missing key additional gRNA dependent features impacting activity.

This variability has made predicting the on-target activity of a given gRNA sequence difficult. While the above results demonstrate the importance of the non-target pool in impacting Cas9 activity, significant future work is needed to understand the specific impact of the number, accessibility (non-target sites in chromatin, for example may not be accessible for Cas9 binding), and similarity of any group of non-target sites. Future studies taking into account the non-target pool could lead to improved models and predictions for the activity of any specific gRNA sequence in a given context and may also enable reductions in off-target cutting, which remains a concern in therapeutic applications of CRISPR technology. Approaches to reduce off target rates have included the use of higher fidelity variants, as discussed above, as well as reducing Cas9 expression levels. Understanding the relative importance of non-target sites and/or utilizing variants with increased PAM specificity may lead to therapies with reduced off-target effects. In addition, some work in the field has focused on engineering Cas9 to recognize a larger number of PAM sites (decreasing PAM specificity) which would open up a larger number of target sites available for therapeutic edits.^15^ However, this would be expected to result in a larger non-target pool and Cas9 variants which have reduced on-target activity i*n vivo*, as recently shown with xCas9 and Cas9-NG.^32,33^ Therefore, in therapeutic applications the opposite approach (increasing PAM specificity) may be expected to improve on-target activity. Future efforts to improve specificity and activity may benefit from engineering enzymes with tighter PAM selection, or leveraging CRISPR nucleases with longer PAM sites, such as the VRER variant of Cas9 or AsCpf1 and LbCpf1.^25,34^ Importantly, slower higher fidelity enzymes can be sped up by increasing PAM specificity. In addition, we anticipate that applications using wild type Cas9, with typically low expression or enzyme levels will be improved by increasing PAM specificity.

Demonstrating that the non-target pool or search space has a real and quantifiable impact on Cas9 activity has the potential to explain a lot of sequence and context dependent variability observed with CRISPR nucleases. However, significant work remains in evaluating how much of this variability can be explained by the non-target pool and how this variation interacts with other important factors impacting Cas9 activity.

## Acknowledgements

We would like to acknowledge the following support: ONR YIP #12043956, and DOE EERE grant #EE0007563. We would also like to acknowledge support from Duke Innovation & Entrepreneurship Initiative. We thank the Duke University School of Medicine for the use of the Sequencing and Genomic Technologies Shared Resource, which provided Next Generation Sequencing service.

## Author contributions

E.A. Moreb and M. Hutmacher performed experiments. E.A. Moreb, M.D. Lynch designed experiments, analyzed results, wrote, revised and edited the manuscript.

