## Supplementary Materials for "CRISPR/Cas “non-target” sites inhibit on-target cutting rates"

### Supplemental Materials

| Table S1: Plasmids and strains used in this study | | | | | | |
|---|---|---|---|---|---|---|
| **Plasmid** | **Insert** | **promoter** | **ori** | **Res** | **Addgene** | **Source** |
| pKDsgRNA-non-targ. | Control gRNA | ptet | pSC101ts | Sm | 89960 | [1] |
| pKDsgRNA-ampG | ampG targeting gRNA | ptet | pSC101ts | Sm | 154923 | This study |
| pKDsgRNA-hisF | hisF targeting gRNA | ptet | pSC101ts | Sm | 154922 | This study |
| pKDsgRNA-malG | malG targeting gRNA | ptet | pSC101ts | Sm | 89951 | [1] |
| pKDsgRNA-mqo | Mqo targeting gRNA | ptet | pSC101ts | Sm | 154921 | This study |
| pKDsgRNA-pgaC | pgaC targeting gRNA | ptet | pSC101ts | Sm | 154919 | This study |
| pKDsgRNA-pheT | pheT targeting gRNA | ptet | pSC101ts | Sm | 154920 | This study |
| pCas9-CR4 | SpCas9 | ptet | p15a | Cm | 62655 | [2] |
| pEM-$Cas9_{D1135E}$ | D1135E | ptet | p15a | Cm | 154931 | This study |
| pEM-Cas9-HF1 | Cas9-HF1 | ptet | p15a | Cm | 89961 | [1] |
| pEM-Cas9-$HF1_{D1135E}$ | Cas9-$HF1_{D1135E}$ | ptet | p15a | Cm | 154924 | This study |
| pEM-eCas9 | eCas9 | ptet | p15a | Cm | 89981 | [1] |
| pEM-$eCas9_{D1135E}$ | $eCas9_{D1135E}$ | ptet | p15a | Cm | 154926 | This study |
| pEM-HypaCas9 | HypaCas9 | ptet | p15a | Cm | 154929 | This study |
| pEM-$HypaCas9_{D1135E}$ | $HypaCas9_{D1135E}$ | ptet | p15a | Cm | 154930 | This study |
| pEM-OptiCas9 | OptiCas9 | ptet | p15a | Cm | 154927 | This study |
| pEM-$OptiCas9_{D1135E}$ | $OptiCas9_{D1135E}$ | ptet | p15a | Cm | 154928 | This study |
| pNT-15 | 15 non-target (NT) sites | -- | pUC | Kan | 154932 | This study |

| | | | | | | |
|---|---|---|---|---|---|---|
| pNT-12 | 12 NT sites | -- | pUC | Kan | 154933 | This study |
| pNT-3 | 3 NT sites | -- | pUC | Kan | 154934 | This study |
| pNT-1 | 1 NT site | -- | pUC | Kan | 154935 | This study |
| Ori- origin of replication, Res - resistance marker, Sm - spectinomycin, Cm- chloramphenicol, Kan - kanamycin, | | | | | | |

| **Table S2: Strains used in this study** | | |
|---|---|---|
| **Strain** | **Genotype** | **Source** |
| BW25113 | F-, λ-, Δ*(araD-araB)567, lacZ4787(del)(::rrnB-3) , rph-1, Δ(rhaD-rhaB)568, hsdR51,* | [3] |
| W (ATCC 9637) | Wild type | [4] |
| NEB - New England BioLabs, Res - resistance marker, Sm - spectinomycin, Cm- chloramphenicol, Kan - kanamycin, Amp - ampicillin | | |

| **Table S2: Oligos & Synthetic DNA used in this study** | | |
|---|---|---|
| **Name** | **Sequence** | **Description** |
| EAM001 | GTGCTCAGTATCTCTATCACTGA | Reverse primer of pKDsgRNA-X KLD reaction |
| EAM002 | ACAGCTACACCCTGGCCGTGgttttagagctag aaatagcaag | Forward primer with EAM001 to make pKDsgRNA-malG |
| EAM003 | CGTGAACGCCACTGGCCGTGgttttagagctag aaatagcaag | Forward primer with EAM001 to make pKDsgRNA-pgaC |
| EAM004 | CCCGCGTTATCTTGGCCGTGgttttagagctaga aatagcaag | Forward primer with EAM001 to make pKDsgRNA-pheT |
| EAM005 | CGCTTCCAGCGATGGCCGTGgttttagagctag | Forward primer with EAM001 to make |

| | | |
|---|---|---|
| | aaatagcaag | pKDsgRNA-hisF |
| EAM006 | TGCTCTTCTCGATGGCCGTGgttttagagctaga aatagcaag | Forward primer with EAM001 to make pKDsgRNA-mqo |
| EAM007 | GCGTCAGCCTGTTGGCCGTGgttttagagctag aaatagcaag | Forward primer with EAM001 to make pKDsgRNA-ampG |
| EAM008 | CAAAACCACCATATTTTTTGGATC | Forward primer for KLD to make D1135E mutation in Cas9 |
| EAM009 | AAAGTCCAACGGTAGCTTATTCA | Reverse primer for KLD to make D1135E mutation in Cas9 |
| EAM010 | Cagacgattcaatagacaataaggtc | Forward primer to make K848A mutation for eCas9 |
| EAM011 | AAGGAAACTTTGTGGAACAATG | Reverse primer to make K848A mutation for eCas9 |
| EAM012 | tctccaaggagtcaattttacca | Amplify Cas9 backbone to insert EAM014 for mutations K1003A and R1060A in eCas9 |
| EAM013 | tcaaagcagttccaacgacg | Amplify Cas9 backbone to insert EAM014 for mutations K1003A and R1060A in eCas9 |
| EAM014 | catgatgcgtatctaaatgccgtcgttggaactgctttgattaagaa atatccagcgcttgaatcggagtttgtctatggtgattataaagttta tgatgttcgtaaaatgattgctaagtctgagcaagaaataggcaaa gcaaccgcaaaatatttcttttactctaatatcatgaacttcttcaaaa cagaaattacacttgcaaatggagagattcgcaaagcgcctctaa tcgaaactaatggggaaactggagaaattgtctgggataaaggg cgagattttgccacagtgcgcaaagtattgtccatgccccaagtc aatattgtcaagaaaacagaagtacagacaggcggattctccaa ggagtcaattttaccaaaaagaaattcggacaa | For NEB HiFi Assembly with PCR amplified by EAM012 and EAM013 to make mutations K1003A and R1060A in eCas9 |
| EAM015 | agcgatcagcgcggcaaacgcGCGATTGGCAAAA CCATCTG | Forward primer to make HypaCas9 mutations (N692A/M694A/Q695A/H698A) |
| EAM016 | GATGATAGTTTGACATTTAAAGAAGAC | Reverse primer to make HypaCas9 mutations |

| | | |
|---|---|---|
| | ATTC | (N692A/M694A/Q695A/H698A) |
| EAM017 | GCGttgtctcgaaaattgattaatggt | Forward primer to make mutation R661A in OptiCas9 |
| EAM018 | tccccaaccagtataacggcgac | Reverse primer to make mutation R661A in OptiCas9 |
| EAM019 | CATcttgaatcggagtttgtctatggtg | Forward primer to make mutation K1003H in OptiCas9 |
| EAM020 | tggatatttcttaatcaaagcagttccaacgac | Reverse primer to make mutation K1003H in OptiCas9 |
| EAM021 | TGAGGCTCGTCCTGAATGATATCAAGCTTGAATTCGTTttctgtggcacgacttctctggactgttcttatcaactccgagggtagactcaatctgaatacttttcgttactggaccctggccgtggggtaaaacggttggcgaactacgatacggaagtaacagtctccctattattctgattggtagcagttaactatcggtctaaccctggccgtgtgggtgggcgtctttgagaactgttgactacacgcttatctgccgaaatagaatactgagtgcctttataaacggttgactaccctggccgtgaggtaagtagatttcttcggaggaataatcgctgagagtcactgaaatcaatagtatcacacctgtttacgactttactgaaccctggccgtgcgggaaacctgttatctaccgattcagaagttcctattgagttactgggagactatcatagttgccagagtgaatccctataccctggccgtgcgggagaactttcagccactatcacggtagaagaatccttttcggagtattttattggtattcacgcaattgccaacagttaccctggccgtgtggttgattcggactgcgatagtaaactgtagataataagaccgctctgacaatacctcaggttttacgactatccagttaaccctggccgtgaggctcagtaaacgaaataatacgataagtagttcccaatctctggttgtctcggtagactcagactaattgttacggacaaccctggccgtggggtacgaagactgattgaggacgcaactacaggttctattttacttattacgccagataggtaaaccagaaatcctcgttaccctggccgtgcggactctgaaaatccgtgttgaataaaggtgtagtagccgttacgcaataccaatctgaatacacttataataccgtgtcaccctggccgtgaggaaacactggattgacttttcaggaagattcgcaactcgtagaagcgtgattactagttacaacgacg | Sequence used in NEB HiFi Assembly with pSMART (Lucigen) to make pNT-15 |

| | | |
|---|---|---|
| | aaagcgtatttaccctggccgtgtggtaagatagtaataaccaaccctgtttccgctgtagtgtaagtgcctttctgaagaattcaatcgtaatctattggcgaaccctggccgtggggagacaataggatttcgctggtttcactctcggaataaagttctcagtagactcaaagacgactttgattatctgagacaccctggccgtgtggatcccacgctgtattatcagattctatcctattcgtagttggaaaagtaagaggcagtgagtaacaagtgcttccaataccctggccgtgaggttatcaactttcaggtaatctctattgtaagaccacgagggacaggcgtattctatagtgacaacttatcagcacgataccctggccgtgcggtaactttctaccctctattgaaggataaggtgttgattggacgattacagtcaggaaccagcctcagtGACGAATTCTCTAGATATCGCTCAATACTGA | |
| EAM022 | ggataaggtgttgattggacga | Forward primer used to delete parts of pNT-15 to make pNT-12, pNT-3, and pNT-1 |
| EAM023 | agtcgtctttgagtctactgaga | Reverse primer used with EAM022 to make pNT-12 |
| EAM024 | tcagtaaagtcgtaaacaggtgt | Reverse primer used with EAM022 to make pNT-3 |
| EAM025 | gactgttacttccgtatcgtagt | Reverse primer used with EAM022 to make pNT-1 |
| EAM026 | AGGCACTTGCTCGTACGACG | Forward primer for amplifying oligo library |
| EAM027 | TTAAGGTGCCGGGCCCACAT | Reverse primer for amplifying oligo library |
| EAM028 | GTGATAGAGATTGACATCCCTATCagtgatagagatactgagcac | Read1 primer |
| EAM029 | GTTGATAACGGACTAGCCTTATTTTAACTTGCTATTTCTAGCTCTAAAAC | Read2 primer |
| EAM030 | GCTAGTCCGTTATCAACTTGAAAAAGTGGCACCGAGTC | Read i5 index |
| EAM031 | ctcatgaccaaaatcccttaacgtgagttttcgttccactga | Read i7 index |

| | | |
|---|---|---|
| EAM032 | AATGATACGGCGACCACCGAGATCTACAC**gacgaact**tcagtggaacgaaaactcacg | Custom i5 barcoded primer (barcode in bold) |
| EAM033 | AATGATACGGCGACCACCGAGATCTACAC**tcggattc**tcagtggaacgaaaactcacg | Custom i5 barcoded primer (barcode in bold) |
| EAM034 | AATGATACGGCGACCACCGAGATCTACAC**caaggtac**tcagtggaacgaaaactcacg | Custom i5 barcoded primer (barcode in bold) |
| EAM035 | AATGATACGGCGACCACCGAGATCTACAC**tcctcatg**tcagtggaacgaaaactcacg | Custom i5 barcoded primer (barcode in bold) |
| EAM036 | AATGATACGGCGACCACCGAGATCTACAC**gtcagtca**tcagtggaacgaaaactcacg | Custom i5 barcoded primer (barcode in bold) |
| EAM037 | AATGATACGGCGACCACCGAGATCTACAC**cgaatacg**tcagtggaacgaaaactcacg | Custom i5 barcoded primer (barcode in bold) |
| EAM038 | AATGATACGGCGACCACCGAGATCTACAC**tctaggag**tcagtggaacgaaaactcacg | Custom i5 barcoded primer (barcode in bold) |
| EAM039 | AATGATACGGCGACCACCGAGATCTACAC**cgtatctc**tcagtggaacgaaaactcacg | Custom i5 barcoded primer (barcode in bold) |
| EAM040 | CAAGCAGAAGACGGCATACGAGAT**acatcg**gactcggtgccactttttcaa | Custom i7 barcoded primer (barcode in bold) |
| EAM041 | CAAGCAGAAGACGGCATACGAGAT**tggtca**gactcggtgccactttttcaa | Custom i7 barcoded primer (barcode in bold) |
| EAM042 | CAAGCAGAAGACGGCATACGAGAT**cactgt**gactcggtgccactttttcaa | Custom i7 barcoded primer (barcode in bold) |
| EAM043 | CAAGCAGAAGACGGCATACGAGAT**attggc**gactcggtgccactttttcaa | Custom i7 barcoded primer (barcode in bold) |
| EAM044 | CAAGCAGAAGACGGCATACGAGAT**gatctg**gactcggtgccactttttcaa | Custom i7 barcoded primer (barcode in bold) |
| EAM045 | CAAGCAGAAGACGGCATACGAGAT**tacaa** | Custom i7 barcoded primer (barcode in bold) |

| | | |
|---|---|---|
| | **g**gactcggtgccactttttcaa | |
| EAM046 | CAAGCAGAAGACGGCATACGAGAT**cgtgat**gactcggtgccactttttcaa | Custom i7 barcoded primer (barcode in bold) |
| EAM047 | CAAGCAGAAGACGGCATACGAGAT**gcctaa**gactcggtgccactttttcaa | Custom i7 barcoded primer (barcode in bold) |
